# LRSim: a Linked Reads Simulator generating insights for better genome partitioning

**DOI:** 10.1101/103549

**Authors:** Ruibang Luo, Fritz J. Sedlazeck, Charlotte A. Darby, Stephen M. Kelly, Michael C. Schatz

## Abstract

**Motivation:** Linked reads are a form of DNA sequencing commercialized by 10X Genomics that uses highly multiplexed barcoding within microdroplets to tag short reads to progenitor molecules. The linked reads, spanning tens to hundreds of kilobases, offer an alternative to long-read sequencing for *de novo* assembly, haplotype phasing and other applications. However, there is no available simulator, making it difficult to measure their capability or develop new informatics tools.

**Results:** Our analysis of 13 real linked read datasets revealed their characteristics of barcodes, molecules and partitions. Based on this, we introduce LRSim that simulates linked reads by emulating the library preparation and sequencing process with fine control of 1) the number of simulated variants; 2) the linked-read characteristics; and 3) the Illumina reads profile. We conclude from the phasing and genome assembly of multiple datasets, recommendations on coverage, fragment length, and partitioning when sequencing human and non-human genome.

**Availability:** LRSIM is under MIT license and is freely available at https://github.com/aquaskyline/LRSIM

## 1 Introduction

Haplotype-resolved or phased genomes are desirable for obtaining insight into diploid or polyploid genomes and studying allele specific expression, allele-specific regulation, and many other important genomic features (Snyder, et al., 2015). However, most of the genomes assembled to date are only a single haploid ‘consensus’ sequence with parental alleles merged arbitrarily (Luo, et al., 2012). Only a few studies have reported true diploid *de novo* assemblies so far. One of the first studies successfully combined Illumina short-read sequencing, PacBio sequencing, and BioNano Genomics genome mapping to published a phased assembly of NA12878 (Pendleton, et al., 2015). The second introduced the FALCON-Unzip algorithm and unraveled three phased genomes, *Arabidopsis thaliana*, *Vitis vinifera* (grape), and the coral fungus *Clavicorona pyxidata*, relying exclusively on PacBio sequencing (Chin, et al., 2016). The third approach generated 605,566 fosmid clones on the YH1 genome and mixed them into 30 clones per pool, each pool containing 0.04% of the diploid genome (Cao, et al., 2015). However, the sample requirements and the associated high cost of the three studies preclude their widespread use.

More recently, Zheng et al. demonstrated a straightforward and less expensive method marketed as GemCode and its successor Chromium by 10X Genomics for creating human diploid *de novo* assemblies (Zheng, et al., 2016). It uses an automated microfluidic system to isolate large DNA molecules in partitions containing sequencing primers and a unique barcode to prepare a library that can be used for Illumina paired-end read sequencing (each partition contains several DNA molecules, which share the same barcode). The library can be generated with as little as 1ng of high molecular weight DNA, far less than alternative approaches. As the sequencing is performed using inexpensive Illumina short-read sequencing, the overall cost is at least two orders of magnitudes lower than pooled fosmid sequencing for a diploid genome and significantly less expensive than current long-read sequencing alternatives.

Simulated sequencing data has proved indispensable for guiding tool development and evaluating tool performance (Earl, et al., 2011). Especially for the complex and unique workflow involved in constructing linked reads, it is essential to develop simulation software that can produce linked reads that capture the most essential characteristics of genome partitioning. To our knowledge, no read simulator for linked reads is available. Thus, we developed LRSim, a <u>L</u>inked <u>R</u>eads <u>Sim</u>ulator, which simulates whole genome sequencing modeled on Chromium Linked Read technology. We integrated all the relevant steps of the Chromium protocol so that it can be used to study linked-read sequencing of different genomes, mutation rates, input libraries, and short-read sequencing conditions *in silico*. We tested LRSim with the 10X Genomics LongRanger variant identification and phasing application and the 10X Genomics SuperNova genome assembler (Weisenfeld, et al., 2016) as well as the independent HapCUT2 phasing algorithm (Edge, et al., 2016) to confirm that alignment, variant identification, phasing, and *de novo* assembly are supported and deliver results similar to those of real data. After studying simulated datasets with multiple parameter combinations, we concluded that 1) the best Phase Block Size (150kb-200kb) can be achieved with a 50x linked-reads depth on a human genome and 1.5M partitions (barcodes); and 2) the standard library preparation protocol tailor-made for the size of humans needs to be adjusted regarding the number of partitions (barcodes) before it can be efficiently used for other genomes of significantly different size, such as *A. thaliana*.

## 2 Methods

### 2.1 Characteristics of real 10X genomic sequences

We analyzed 13 publicly available real datasets processed by Chromium’s LongRanger analysis pipeline to derive models and characteristics (see Supplementary Note). We identified 1.5 million partitions with more than 100 reads each in NA12878 (Figure 1). In Chromium’s 16bp barcode we observed 2.09% total errors, which is slightly higher than the 1.78% estimated by base quality (Supplementary Table 1). Interestingly, two of the barcodes were found to be consistently over-represented in all 13 samples

**Figure 1.**
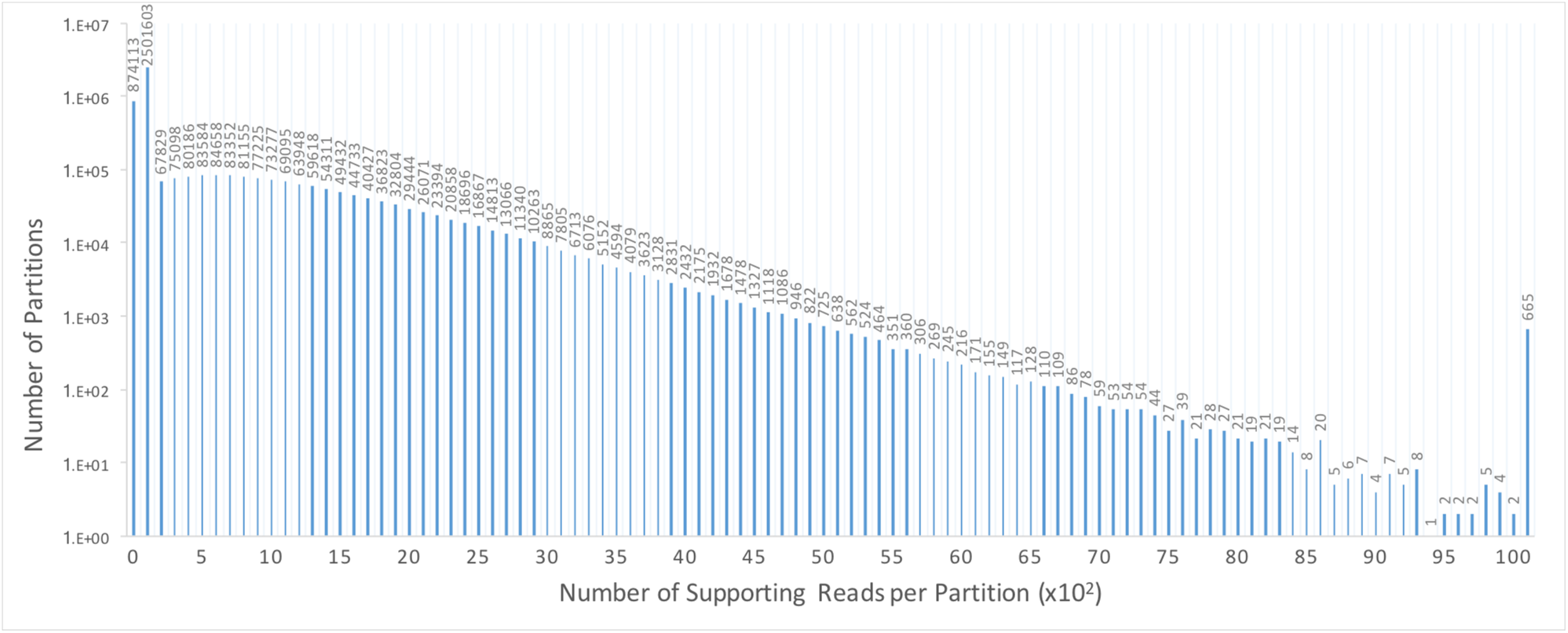
The distribution of number of supporting reads per partitions. About 1.5 million partitions are supported by more than 100 reads.

**Table 1.**
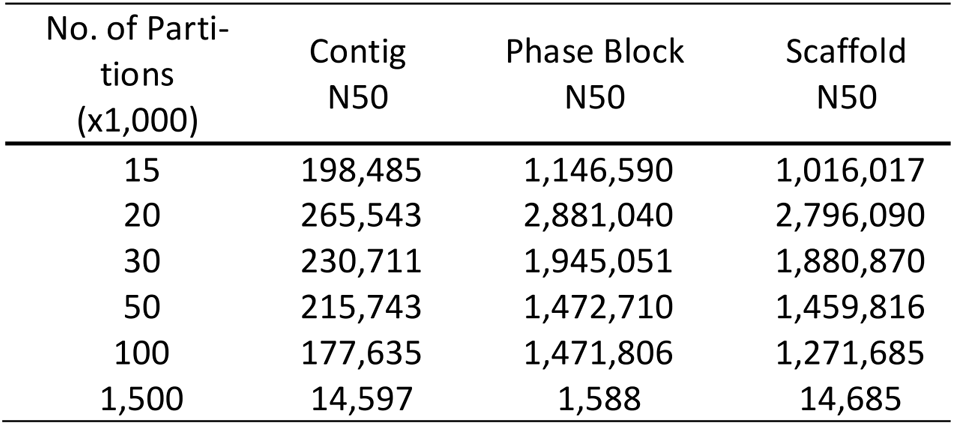
The Contig N50, Phase Block N50 and Scaffold N50 of the *A. thaliana* genome with 6 different partition numbers.

(“GTATCTTCAGATCTGT,” “GTGCCTTCAGATCTGT,” Supplementary Table 2). Considering all regions ≥50 base-pairs that had zero coverage, we found that the Chromium 80x NA12878 dataset left 164Mb (5.47%) uncovered by any read alignment, versus 209Mb (6.97%) that had zero coverage in the NIST 300x NA12878 Illumina only paired-end read dataset (Supplementary Table 3). In the NA12878 sample specifically, in average 10 DNA molecules on average were allocated to each partition (Figure 2). The weighted molecule length peaked at around 40-50kbp (Figure 3). The distribution of molecule coverage peaked at 0.2x coverage (Figure 4). Reads were generally uniformly distributed along the genome, as well as along the molecule, although surprisingly, we observed chromosome 21 with unexpectedly high depth in all samples (Figure 5; Detailed views in Supplementary Figure 1a, b).

**Figure 2.**
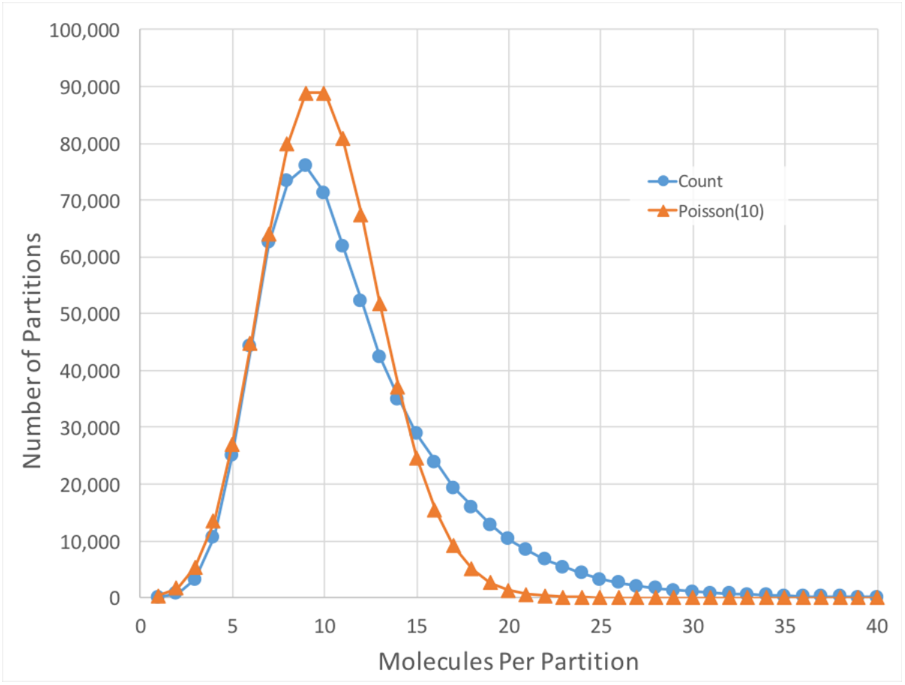
Distribution of the number of molecules per partition for NA12878.

**Figure 3.**
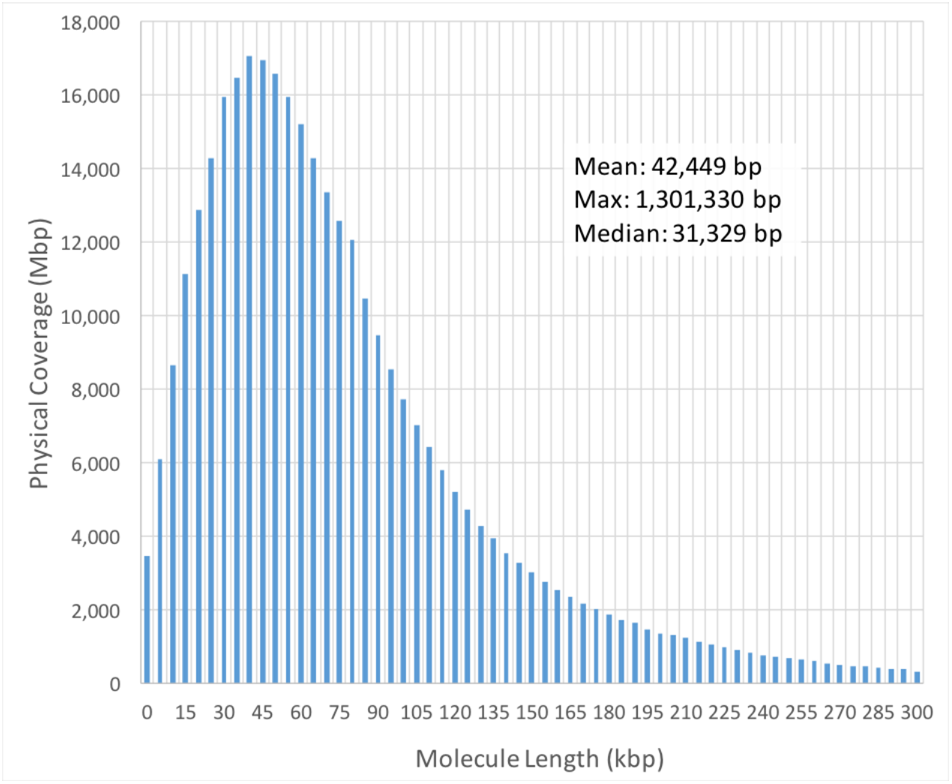
Weighted molecule length distribution for NA12878. Physical Coverage equals 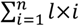, where l is the length of the molecule and n is the number of molecules in that size range.

**Figure 4.**
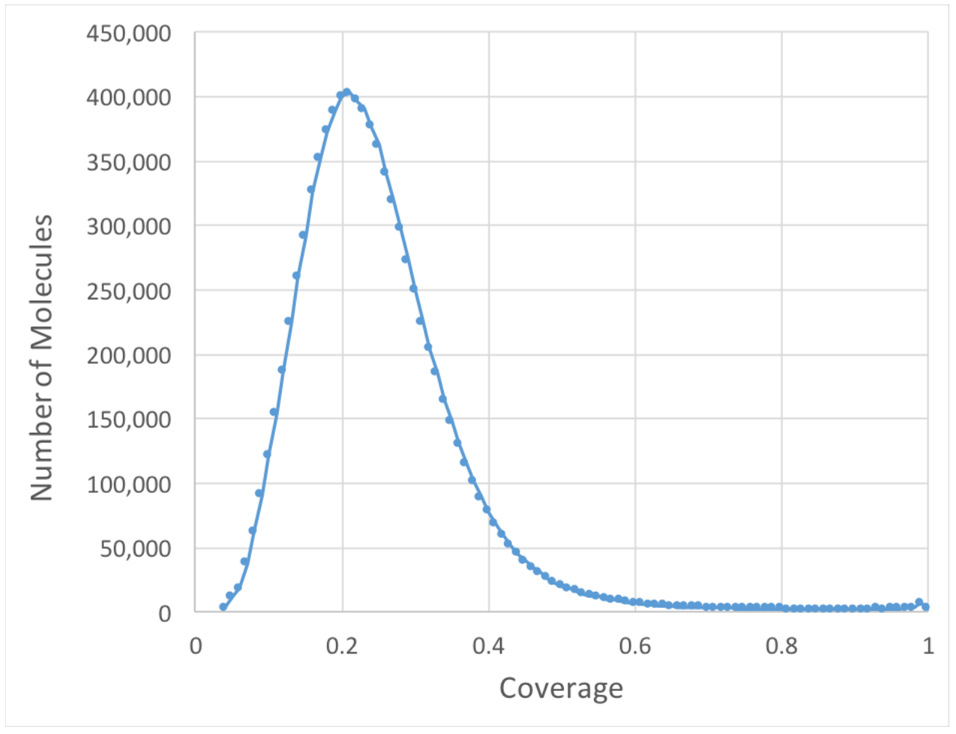
Distribution of molecule coverage for NA12878.

**Figure 5.**
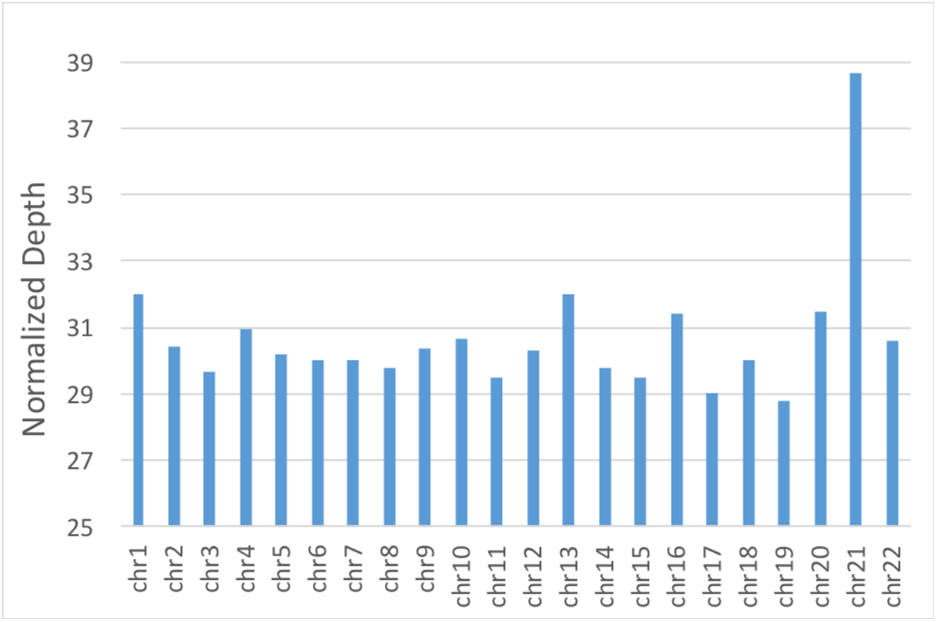
Average depth of 13 samples per chromosome. The depths were normalized to the sample with the lowest average depth (NA24149, 30.36x).

**Table 2.** Contig N50, Phase Block N50 and Scaffold N50 of the *A. thaliana* genome with 4 different combinations of number of read pairs and number of partitions.

| No. of read pairs (M) | No. of Partitions (x1,000) | Contig N50 | Phase Block N50 | Scaffold N50 |
| --- | --- | --- | --- | --- |
| 9 (16-fold) | 20 | 233,233 | 1,027,768 | 899,826 |
| 18 (32-fold) | 20 | 265,543 | 2,881,040 | 2,796,090 |
| 27 (48-fold) | 20 | 221,680 | 1,971,701 | 1,896,517 |
| 27 (48-fold) | 30 | 241,319 | 1,979,723 | 1,688,453 |

**Table 3.** Contig N50, Phase Block N50 and Scaffold N50 of the *A. thaliana* genome with 6 different molecule numbers per partition.

| No. of Molecules per Partition | Contig N50 | Phase Block N50 | Scaffold N50 |
| --- | --- | --- | --- |
| 1 | 46,993 | 72,400 | 54,769 |
| 4 | 249,371 | 2,105,517 | 2,020,687 |
| 7 | 274,133 | 1,869,856 | 2,074,127 |
| 10 | 265,543 | 2,881,040 | 2,796,090 |
| 15 | 232,708 | 2,860,223 | 2,416,675 |
| 20 | 245,894 | 1,920,175 | 1,878,113 |

### 2.2 Simulator design and performance

Following our observations from the 13 real datasets, we included the following parameters in our simulator: *x*: number of read pairs; *t*: number of partitions; *f*: mean molecule length; and *m*: mean number of molecules per partition. The Poisson distribution is used to sample data from *f* and *m* by default; it can easily be switched to other functions. Figure 6 shows the overview of the LRSim workflow. Briefly, LRSim first uses SURVIVOR (Jeffares, et al., 2017) to simulate homozygous and heterozygous SNPs, indels and structural variants within the user-specified genome sequence. Second, LRSim uses DWGSIM (GitHub: nh13/DWGSIM) to mimic the error profile of Illumina paired-end reads and generates 50% more reads than the user requested. Third, considering the Illumina reads as a pool, LRSim emulates the process of linked-read sequencing by attaching barcodes to reads selected from the pool. For a human genome using default parameters, the memory consumption peaks at 48GB, and starting from scratch takes about five hours to finish using 8 threads, or 1.5 hours if only the linked reads related parameters *x*, *f*, *t* or *m* are altered, thus avoiding rerunning the expensive DWGSIM stage.

**Figure 6.**
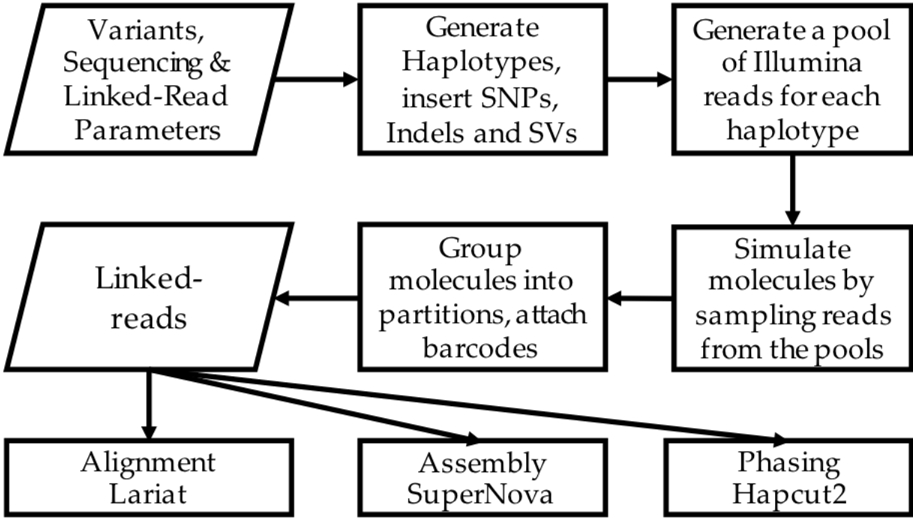
LRSim workflow.

## 3 Results

### 3.1 Effect on molecule size (*f*)

One of the critical requirements of linked read construction is extracting high-quality, high-molecular weight DNA from the sample. To study how the molecule size changes the performance of linked-read sequencing, using human reference genome GRCh38, we simulated six datasets of different molecule sizes (*f*: 20, 50, 100, 150, 200 and 250kbp), with 600 million read pairs (*x*), 1.5 million partitions (*t*) and 10 molecules per partition (*m*). Instead of simulating random variants, which may not mimic the characteristics of real variants, we used 3.2M phased SNPs and indels from NA12878 (Supplementary Note). The datasets were processed by LongRanger and phased by HapCUT2 (Edge, et al., 2016) using a 48- core Intel E7-8857 v2 @3GHz machine with 1TB memory, running on average 1.5 days each. The sum of bases of different phase block sizes for six simulated datasets and NA12878 (Supplementary Note) are shown in Figure 7. The results show that the NA12878’s performance lies between molecule size 50kbp and 100kbp, corroborating our observation in the weighted molecule length distribution of NA 12878 (Figure 4). The divergence between 50kbp and NA12878 can be explained by the fact that the real data is platykurtic and has a longer tail on long molecules, which is an outcome that highly depends on the quality and length of DNA input.

**Figure 7.**
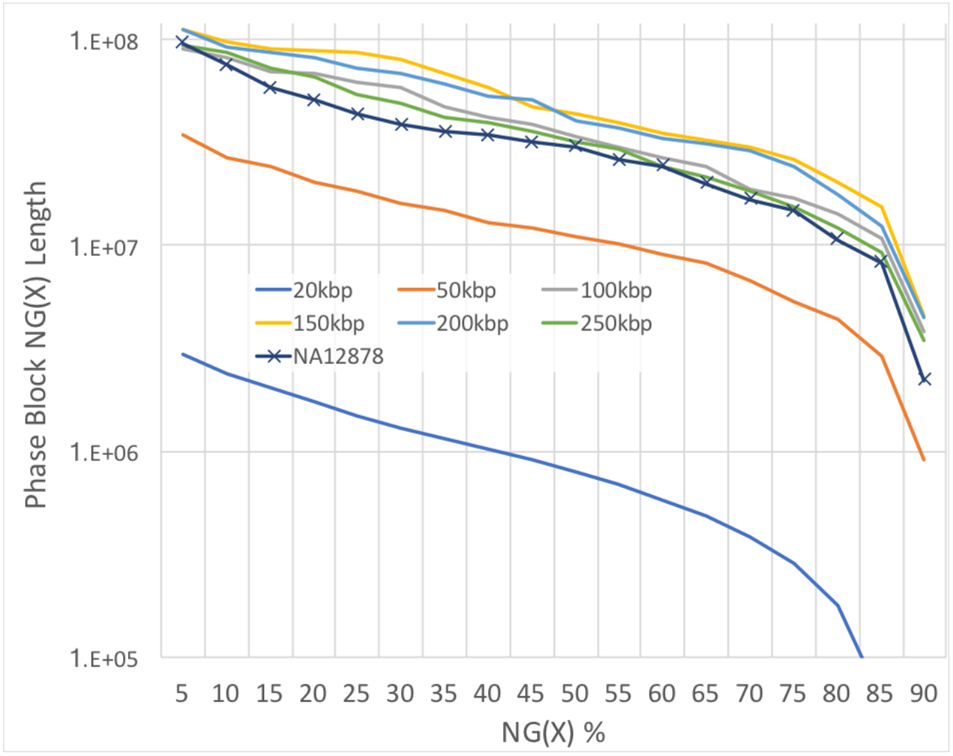
NG graph showing an overview of phased block sizes of 7 datasets. NG(X) is defined as X% of the genome is in phased blocks equals to or larger than the NG(X) length.

We further noted that the phase block N50 sizes did not monotonically increase with longer molecules and instead plateaued at 200kbp. On investigation, we determined the cause to be that given a constant number of total reads, the coverage per molecule decreases proportionally to the molecule size. For example, if we increase the molecule size from 50kbp to 250kbp, the coverage decreases from ~0.2x to ~0.04x, and the average distance and standard deviation of the distances between reads rises, thus leading to shorter phase blocks.

### 3.2 Effect on the number of partitions (*t*)

We simulated six datasets with varying numbers of partitions (*t*), including 15k, 20k, 30k, 50k, 100k and the standard1.5 million, to study the impact of the number of partitions on the assembly using the *A. thaliana* genome (TAIR10).

We kept the other parameters constant at 18 million read pairs (*x*), 50 kbp mean molecule size (*f*) and 10 molecules per partition (*m*). Using the same computer, SuperNova finished each assembly within 2 hours. The best assembly, as measured by contig N50, scaffold N50 or phase block N50, was from the dataset with *t*=20,000 partitions (Table 1). Intriguingly, using 1.5 million partitions, which is the default for the 10X Chromium platform, the results are orders of magnitude worse for this reduced genome size. The deficiency in the assembly performance can be attributed to insufficient coverage per molecule. For a 3Gbp genome with the default parameters, the coverage per molecule is 0.2x on average. For *A. thaliana*, where the genome size is 20 times smaller (150Mbp vs. 3Gbp), if all parameters remain in the default mode, the number of reads allotted to each molecule will be 20 times smaller, i.e. 0.01 coverage. The excessively low coverage increases the mean distance and its standard deviation between reads, which confounds the genome assembler; it also largely removes the chance for reads belonging to the same molecule to cover multiple heterozygous variants, which is essential for phasing. Therefore, we suggest adjusting the number of partitions according to the genome size, or increasing the overall sequencing coverage and subsample the partitions proportionally. This will result in sufficient coverage per partition, which improves the assembly results using linked reads.

### 3.3 Effect on coverage (*x*)

Genome assembly using Illumina short reads requires careful control of the sequencing coverage. Shallow coverage decreases the maximum usable kmer-size (to achieve the minimum requirement for kmer depth), thus limiting the ability to disentangle repetitive sequences. Excessively deep coverage leads to a lower signal-to-noise ratio because of the saturation of authentic information and the accumulation of more random errors in the ‘assembly graph’, thus decreasing the performance of the assembly outcome (Li, et al., 2012; Zerbino and Birney, 2008). The best practice for sequencing depth ranges from around 30x to 100x coverage, depending on affordability (a longer kmer-size can be used with greater depth, but this is limited by the length of read input and read errors) and the genomic nature of different species, including their heterozygosity, heterogeneity and repetitiveness. It is less clear how the coverage of linked reads changes the performance of genome assembly.

Using the *A. thaliana* genome, we simulated four datasets with three different numbers of read pairs (*x*), 9, 18 and 27 million, which equates to 16-, 32- and 48-fold of the genome, respectively, and two different numbers of partitions, 20,000 and 30,000 for *x*=27. The molecule length (*f*) was held constant at 50kbp, as was the number of molecules per partition at *m*=10. We used SuperNova to assemble the four datasets. The results are shown in Table 2. We found that 18 million read pairs (32-fold) with 20,000 partitions achieved the best Contig N50, Phase Block N50 and Scaffold N50. Interestingly, the assembly result of 27 million read pairs (48-fold) was worse than 18 million on all three metrics, and only improved slightly on Contig N50 and Phase Block N50 after increasing the number of partitions to 30,000 (to keep the molecule coverage the same as 18 million read pairs). This indicates that the sequencing coverage itself rather than the molecule coverage makes a difference in linked-reads genome assembly using the SuperNova assembler.

### 3.4 Effect on the number of molecules per partition (*m*)

The number of molecules per partition is usually determined by sample preparation technologies and cannot be easily modified except by carefully controlling the total amount of input DNA. Thus, the number needs to be carefully selected and verified before production. A lower number of molecules per partition requires a larger number of barcodes to arrive at the same number of molecules. A higher number of molecules per partition requires fewer barcodes, but increases the chance of two molecules coming from the two haplotypes in the same genome position. Given the number of barcodes, the number of molecules per partition will increase or decrease the coverage per molecule, which changes the performance of genome assembly and phasing.

Using the *A. thaliana* genome, we simulated 6 datasets with 1, 4, 7, 10, 15 and 20 molecules per partition, with the number of read pairs (*x*=18 million), molecule length (*f=*50,000) and number of partitions (*t*=20,000). The assembly results are shown in Table 3. Phase Block N50 and Scaffold N50 peaked with 10 molecules per partition, while contig N50 peaked with 7. The metrics went down significantly with only 1 molecule per partition; the reason remains unclear. We note that molecule coverage increased to 2 with only 1 molecule per partition; this could confound phasing algorithms oblivious to conflicting alleles caused by sequencing error within a molecule. Also, genomic regions being covered decreases with the same number of partitions but less molecules per partition.

## 4 Discussion

In this paper, we presented an analysis of 13 real datasets of linked reads generated by 10x Genomics technology. We implemented a linked-read simulator named LRSim to allow fine tuning of both the type and number of variants and Illumina read specifications, and full control of important parameters for linked-reads we identified in real datasets, including 1) the number of read pairs; 2) the number of partitions; 3) the mean molecule length; and 4) the mean number of molecules per partition. The performance of the simulated data matched real dataset NA12878 closely in phasing. We concluded that 1) from the phasing results of 6 simulated datasets with different mean molecule lengths and a real dataset of NA12878 that if constrained at a certain sequencing depth, the best molecule size to achieve the best phase block size needs to be meticulously chosen. This can be done by wet-lab experiments, but would be more efficient with a simulator in silico; 2) experiments on 6 simulated A. thaliana datasets with a different number of partitions demonstrated a substantial degradation in assembly performance with an improper number of partitions, which leads to insufficient coverage per molecule; 3) an appropriate sequencing depth needs to be chosen for different applications and species before sequencing to achieve the best performance out of linked-reads.

In our study, linked reads enabled much longer contigs, scaffolds and phase blocks on both the human genome and *A. thaliana* than using Illumina short-reads for genome assembly. The better outcomes, in turn, broaden the horizons for studies of allele specific expression, allele-specific regulation, and many other important genomic features critical to precision medicine. Furthermore, numerous other sequencing applications, such as metagenomics and RNA-seq, could potentially benefit from linked-read data. Linked-read technology is promising, and we believe that more complex genomics workflows will include and benefit from it. We therefore encourage users to use LRSim to aid in the development of these new workflows.

## Acknowledgements

We thank Steven Salzberg for his valuable comments and editing the manuscript. We also thank the team of 2016 CSHL-NCBI Hackathon for helpful discussions. We would also like to thank Deanna Church, David Jaffe, and Patrick Marks from 10X Genomics for their helpful discussions during the development of LRSim.

